# Vis-OCT Explorer: an open-source software for visible-light optical coherence tomography data processing

**DOI:** 10.1101/2025.10.01.679626

**Authors:** Weijia Fan, Fengyuanshan Xu, Roman Kuranov, Ronald Zambrano, Jacqueline Chen, Jiahui Wu, Seung Hyen Lee, Kenny Q. Trang, Rukhsana Mirza, Shira Simon, Fabio Lavinsky, Xiaorong Liu, Jeffrey L. Goldberg, Alex S. Huang, Joel S. Schuman, Hao F. Zhang

## Abstract

**Background and objectives:** Visible-light optical coherence tomography (vis-OCT) has enabled the visualization of retinal structures and functions beyond the capabilities of conventional OCTs. However, to reconstruct high-quality images, vis-OCT requires special post-processing, including balanced detection. An open-source, standardized vis-OCT data processing software is essential for clinical applications and translation of vis-OCT.

**Methods:** We developed Vis-OCT Explorer, an open-source, modular Python-based software for processing vis-OCT images. In addition to the standard spectral-domain OCT processing pipeline – including k-space resampling, dispersion compensation, and fast Fourier transformation – Vis-OCT Explorer offers unique dual-spectrometer balanced detection, short-time-Fourier transformation (STFT) based dispersion compensation coefficient optimization, and GPU-accelerated processing. We evaluated the reconstruction performance by quantifying a quality index extracted from individual B-scan images. We also assessed the repeatability of retinal thickness measurements by five operators on images acquired from different testing sites using the intraclass correlation coefficient (ICC) analysis.

**Results:** Balanced detection and STFT-based dispersion compensation significantly increased the quality index of reconstructed B-scan images. ICC values of the retinal nerve fiber layer (RNFL) and ganglion cell-inner plexiform layer (GCIPL) thickness measurements from four testing sites exceeded 0.8 in 87.5% of the macular-centered images. The ICC of RNFL thickness measurements on all optic nerve head-centered images is above 0.8, showing strong repeatability across users.

**Conclusions:** Vis-OCT Explorer provides high-quality image processing and enables highly repeatable measurements on vis-OCT human retinal images. It facilitates future multicenter clinical tests to validate vis-OCT’s clinical efficacy.

## Introduction

Optical coherence tomography (OCT) is a noninvasive imaging technique that generates high-resolution three-dimensional (3D) images. Despite the widespread use of OCT in pre-clinical investigations and human disease diagnosis [1], the availability and processing of OCT raw fringe data have long been recognized as critical factors in advancing OCT data standardization [2]. However, the lack of accessible tools to process OCT data has hindered broader efforts to standardize across the field.

In the past few years, multiple open-source software applications for OCT have been published, facilitating the broadening of the OCT research community. For example, Vortex is a C++ library with Python bindings for OCT hardware control and real-time image visualization [3]. OCTSharp offers a comprehensive OCT imaging solution built on .NET C#, integrating hardware control and image acquisition [4]. OCTProz is another real-time OCT image processing software that allows customized or virtual hardware binding [5]. However, most software applications require appropriate hardware setups to operate, which are not always available or standardized among researchers. Additionally, these tools prioritize real-time processing speed over post-processing optimization, and common image analyses used by clinical researchers are often unavailable.

Advanced post-processing of the interferogram data has been shown to significantly enhance OCT image quality, particularly for visible-light OCT (vis-OCT), whose theoretically high axial resolution may be compromised without appropriate image reconstruction [6, 7]. Methods such as balanced detection, dispersion compensation, and image registration have been developed to improve signal-to-noise ratio (SNR), sharpness, and overall image quality for vis-OCT [6, 8, 9]. With refined image processing, finer-scale analysis of human retinal images, such as quantitative measurement of inner plexiform layer (IPL) sublayers [10] and morphometric analysis of the retinal pigment epithelium (RPE) and Bruch’s membrane (BM) [11], becomes feasible.

To the best of our knowledge, an open-source OCT post-processing tool consisting of balanced detection, optimized reconstruction for OCT and OCTA, and retina image thickness measurements remains unavailable. We developed, tested, and validated the open-source Vis-OCT Explorer in Python to fill this void. We report on the architecture, key algorithms, and hardware requirements for operating Vis-OCT Explorer. We also present a repeatability test across five institutions on the performance of image reconstruction and manual retinal thickness measurement using Vis-OCT Explorer.

## Methods

### 2.1 Software solutions and hardware requirements

We used Python (version 3.9) to develop Vis-OCT Explorer. Specifically, we used PySide6 [12], a Python module for Qt, for the graphical user interface (GUI) development, and PyQtGraph [13] for displaying images and graphs. Figure 1 shows a representative screenshot of the GUI.

**Figure 1.**
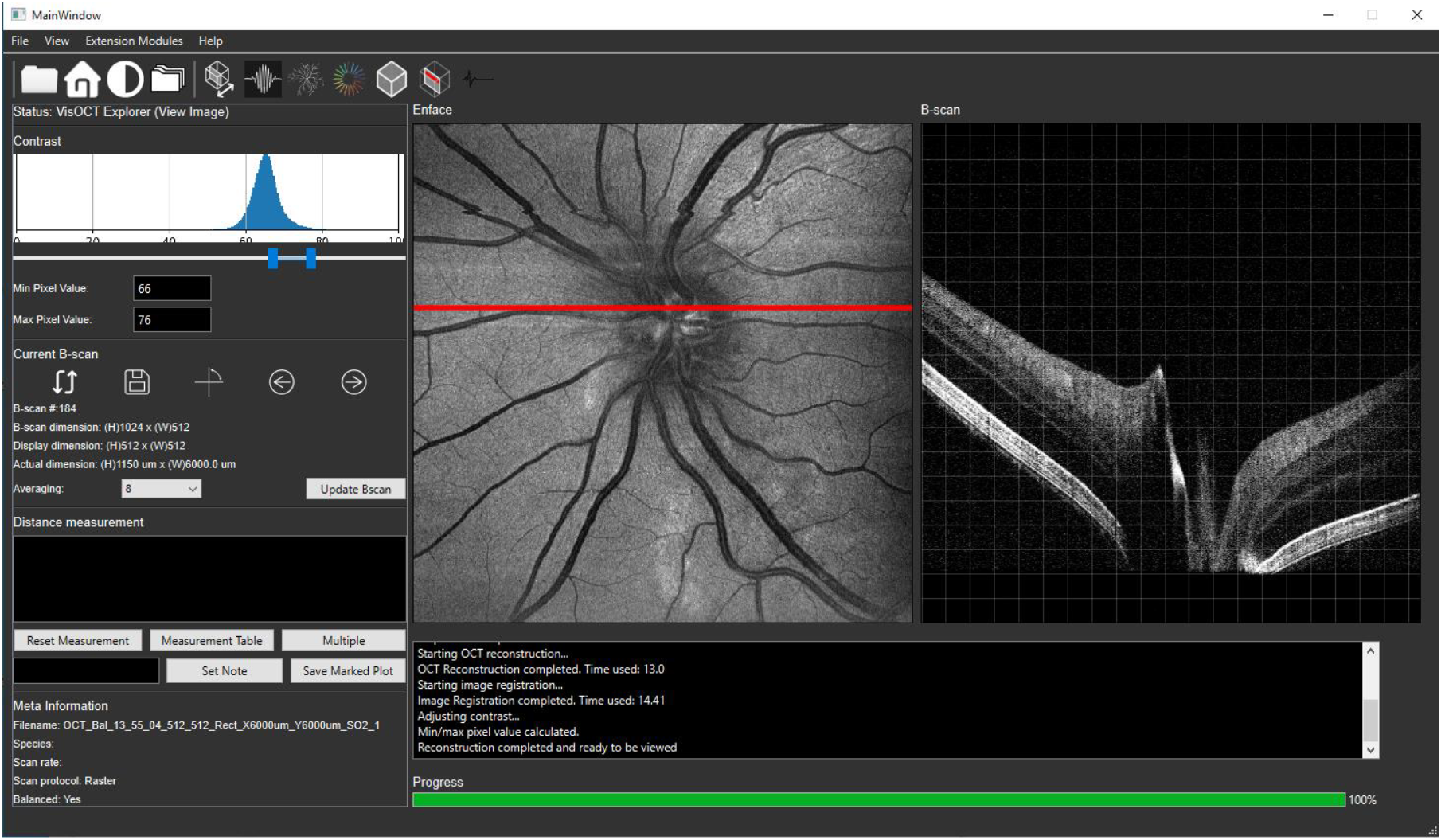
A screenshot of the Vis-OCT Explorer GUI.

Vis-OCT Explorer supports both CPU and GPU processing. We used GPUtil to check the GPU availability and ensure that the user’s computer meets the minimum requirement of 4 GB GPU VRAM. Signal processing and image analysis functions were mainly implemented using the SciPy [14] and NumPy [15] packages for CPU-based processing and the CuPy [16] package for GPU-based processing. The methods and results presented in this work are based on the GPU version. The entire package, along with the open-source code, is available from the GitHub repository at github.com/FOIL-NU/VisOCT-Explorer.

### 2.2 Vis-OCT image preprocessing and reconstruction pipeline

Figure 2 shows the GPU-based image reconstruction pipeline. The GUI was handled on the CPU, and the core back-end processing was performed on the GPU. The data was transferred between DRAM and GPU VRAM in batches for efficient processing. Icons on the GUI are labeled with their corresponding functions. Vis-OCT Explorer provides solutions to reconstruct structural OCT volumes and OCT angiography (OCTA) volumes from spectrometer interferogram data (hereafter referred to as raw data). The software accepts either a single spectrometer dataset for direct reconstruction or a pair of spectrometer datasets for balanced detection reconstruction, where the two datasets should be π phase shifted for noise cancellation [9]. In addition to the raw data, users must supply a wavelength-to-pixel map [12] for the spectrometer. For balanced detection processing, an inter-spectrometer calibration map is also required [13, 14].

**Figure 2.**
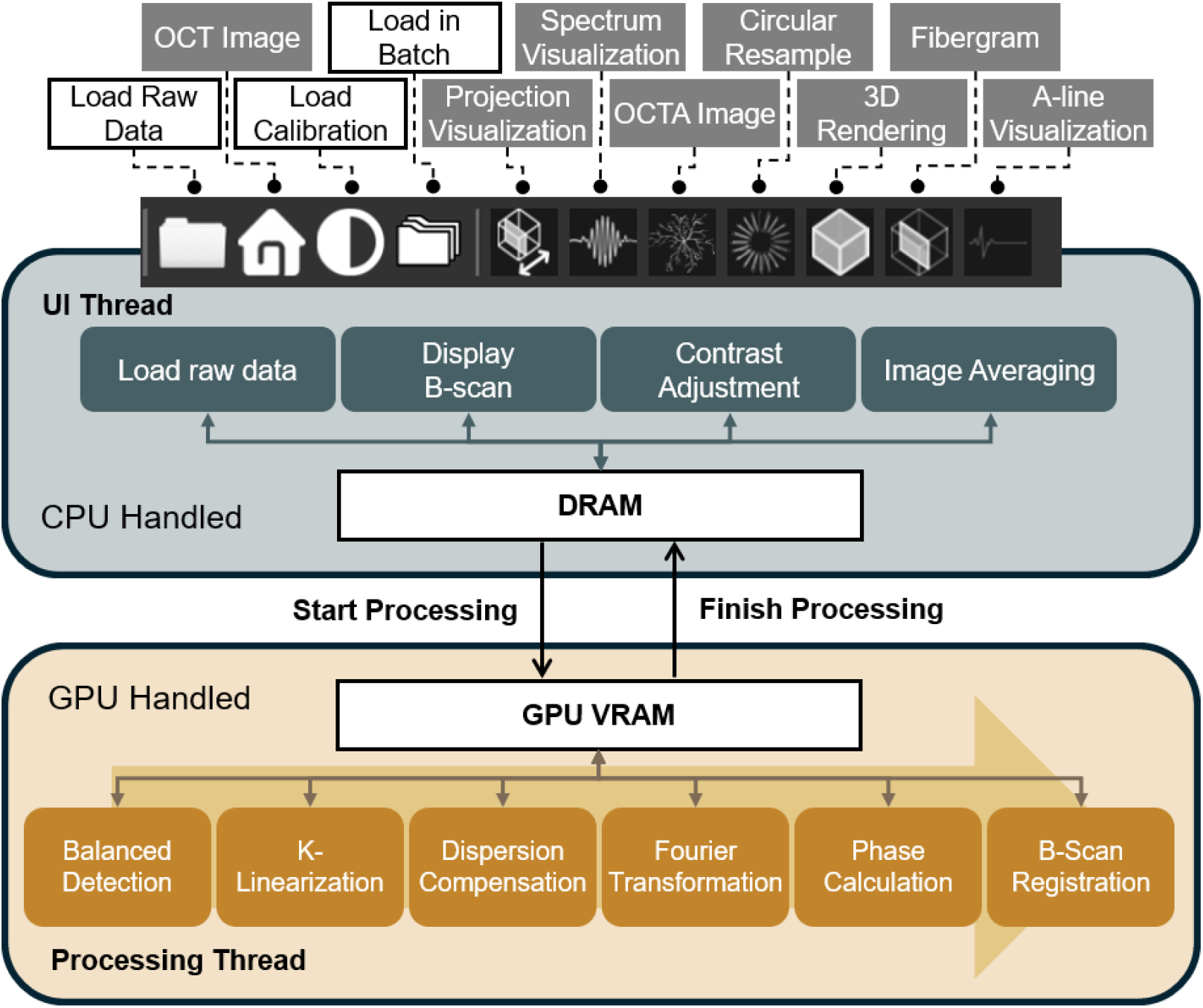
Schematic of GPU-based image reconstruction.

The uploaded raw data underwent the standard spectral-domain OCT reconstruction pipeline, including background subtraction, wavenumber (k) space resampling, dispersion compensation, and fast Fourier transformation (FFT) [2]. For a single spectrometer input, the mean intensity across all spectra was used as the reference arm intensity, or background signal, and subtracted from the input spectra to obtain the interferogram signal. For balanced detection from two spectrometer datasets, one dataset was interpolated using a non-linear spline method to align with the other one based on the inter-spectrometer calibration map [9]. Then, both spectrometer datasets were normalized by their respective background signal intensities before being subtracted from each other. During this process, the non-interferometric background signals were removed, and the interferometric signals were doubled due to the π phase-shifted relationship.

After background removal, we resampled the raw data to a linear wavenumber (k) space using linear interpolation. We then applied a Tukey window for spectral shaping to suppress the side lobe effect after FFT [17]. Next, we digitally corrected the chromatic dispersion mismatch between the reference and sample arms. To automatically estimate 2^nd^- and 3^rd^-order dispersion compensation coefficients, we used short-time Fourier transformation (STFT) to reconstruct sub-band OCT images; then, we optimized the dispersion compensation coefficients by minimizing the axial discrepancy across all the sub-band structural images [18]. The optimized coefficients were applied to calculate the phase correction and added to the original fringes.

Following preprocessing, FFT converted the Fourier space spectral data into interpretable OCT structural images. If the OCTA scan protocol was used with repeated B-scan data, the complex values from the FFT were used to extract phase fluctuation for blood flow signals based on a combined optical microangiography (OMAG) algorithm [19] and split-spectrum analysis [20]. Top-hat filters were then applied to extract blood vessels from the background. After all the volumes were reconstructed, we used the discrete Fourier transform (DFT) to align the structural OCT B-scans at the sub-pixel level and correct for possible motions [21]. The alignment correction was also applied to the OCTA B-scan volumes.

When a CUDA-compatible GPU was available, the GPU handled all the FFT and non-linear interpolation-related operations. Specifically, the spectrometer raw data was transferred from RAM to GPU VRAM in chunks. Balanced detection, dispersion compensation coefficient optimization, FFT, and B-scan registration were key operations implemented using GPU-based functions, significantly reducing post-processing time.

### 2.3 Vis-OCT image post-processing and analysis

After image reconstruction, Vis-OCT Explorer automatically saves the image volumes in .npy format to the local folder containing the corresponding spectrometer raw data. Following biomedical image data interchange conventions [22] and to ensure compatibility with external analysis tools, users can export the reconstructed volumes in .dicom or stacked .tiff formats.

Additional post-analysis functions, including contrast adjustment, registration and averaging, and digital resampling [23], are available in Vis-OCT Explorer. By default, the reconstructed images were displayed using a logarithmic intensity scale for visualization. The grayscale intensity map’s lower and upper limits were empirically set based on the mean value and standard deviation of image volume intensity. A histogram of the image volume intensities was also provided, allowing users to interactively adjust the contrast and brightness of the displayed B-scan and en-face images.

Vis-OCT Explorer offers two image registration and averaging modes. One is the averaging of multiple volumes using a temporal speckle averaging scanning protocol [24]. The repeated volumes often suffer from motion artifacts when multiple volumes are acquired from the same retinal region sequentially. To correct this, we performed 2D registration based on blood vessel features in the OCTA mean intensity projection *en-face* image to generate a deformation field, which we then applied to each structural OCT *en-face* image to remove bulk motion [18]. The B-scan images from each deformed volume were further registered before averaging. Alternatively, users can average neighboring B-scans within a single high-density volume. After specifying the number of B-scans to average, we registered the specified number of adjacent B-scans based on autocorrelation to correct minor motion artifacts before averaging and saving the result.

Furthermore, retinal images of the optic nerve head (ONH) typically require measurements from a circular B-scan with a specific radius centered at the ONH. To support this clinical practice, when data from direct circular scans is not available, Vis-OCT Explorer provides a function to resample image volumes into pseudo-circular B-scans digitally [23]. Given that the ONH is not always at the center of the field of view, we allow users to specify the center of the circle as the ONH manually and set the radius parameter as desired. A-lines along the circumference of the resampling circle were automatically selected and grouped with 12 neighboring A-lines in the radial direction. These A-lines were then registered based on autocorrelation and averaged to form one representative A-line on the pseudo-circular B-scan [23]. This digital resampling and averaging create a circular scan surrounding the ONH from raster scan volumes for retinal layer measurements.

### 2.4 Animal ethics and human imaging

We tested Vis-OCT Explorer on human images and small animal images, including those of mice and tree shrews. We strictly followed the institutional review board (IRB) approved protocols for human imaging at Northwestern University, the University of Virginia, Wills Eye Hospital, Stanford University, and the University of California, San Diego, using vis-OCT systems installed at these institutions.

Small animal images were acquired using multiple repeated volumes to enable speckle noise reduction and OCTA processing. All animal handling procedures followed protocols approved by the Institutional Animal Care and Use Committees (IACUC) at Northwestern University and the University of Virginia. Mice were imaged at both Northwestern University and the University of Virginia, and tree shrews were imaged only at the University of Virginia. Mice were anesthetized using a ketamine and xylazine cocktail injection, while tree shrews were anesthetized with 5% isoflurane delivered with supplemental oxygen. For both species, the eyes were dilated with 1% tropicamide and kept moist with artificial tears. During imaging, animals were stabilized in custom-built holders designed for each species.

### 2.5 Quantitative metrics for software performance validation

We evaluated the performance of Vis-OCT Explorer using eight human volumetric datasets acquired with raster scanning patterns centered on either the optic nerve head (ONH; four datasets) or the macula (four datasets), obtained from five different research institutes as mentioned in Section 2.4. Six datasets were acquired using balanced-detection dual spectrometer systems [9], and two were acquired using a single spectrometer system. In addition, we included three mouse and three tree shrew datasets, each containing 3–5 repeated volumes and two repeated B-scans. These small-animal datasets were used to evaluate the software’s capability for OCTA reconstruction and efficient inter-volume image registration.

We assessed the image reconstruction quality by comparing Vis-OCT Explorer reconstructions with images generated by traditional OCT reconstruction methods previously reported in open-source real-time OCT acquisition software [5]. The most time-consuming processing steps in Vis-OCT Explorer include refined spectral matching via non-linear B-spline interpolation for balanced detection, and dispersion compensation coefficient optimization using sub-band image registration. In contrast, conventional methods typically do not use balanced detection; if they do, spectral matching is performed using hardware-based balancing or linear interpolation. Dispersion compensation is typically implemented by manually entering coefficients (see Table 1). These simplifications enable real-time image reconstruction but may compromise reconstruction quality in some situations.

**Table 1.** Comparison between raw data preprocessing methods of the traditional OCT reconstruction pipeline and the Vis-OCT Explorer reconstruction pipeline.

|  | Traditional OCT Reconstruction | Vis-OCT Explorer OCT Reconstruction |
| --- | --- | --- |
| Balanced Detection | Non or linear interpolation | B-spline interpolation |
| K-linearization | User input resampling curve | User input resampling map |
| Dispersion Compensation | Manually selected coefficient | STFT based coefficient optimization |

To quantitatively compare image quality between the two processing pipelines, we computed a quality index (QI) for each B-scan from six human datasets acquired with balanced detection. Following the established method [16], we calculated the quality index (QI) based on the 1^st^, 75^th^, and 99^th^ percentile values of the B-scan intensity histogram, corresponding to low signal amplitude, background noise, and saturation levels, respectively. This QI quantifies not only the signal-to-noise ratio, but also the contrast between tissue structures with varying reflectivity.

Next, we validated the repeatability of Vis-OCT Explorer in measuring retinal anatomical features by having five operators analyze these eight human datasets. For macular-centered images, one operator randomly selected five B-scans per dataset and marked four locations per B-scan to measure the RNFL, IPL, and GCL. The other four operators repeated these measurements at the exact locations using a mesh grid overlay for reference, as shown in Figure 3(a). Each B-scan was generated by averaging 16 adjacent frames to reduce speckle noise. For ONH-centered scans, all five operators measured RNFL thickness at every 30-degree interval along a digitally resampled B-scan generated at a 1.5 mm radius from the ONH center, as shown in Figure 3(b).

**Figure 3.**
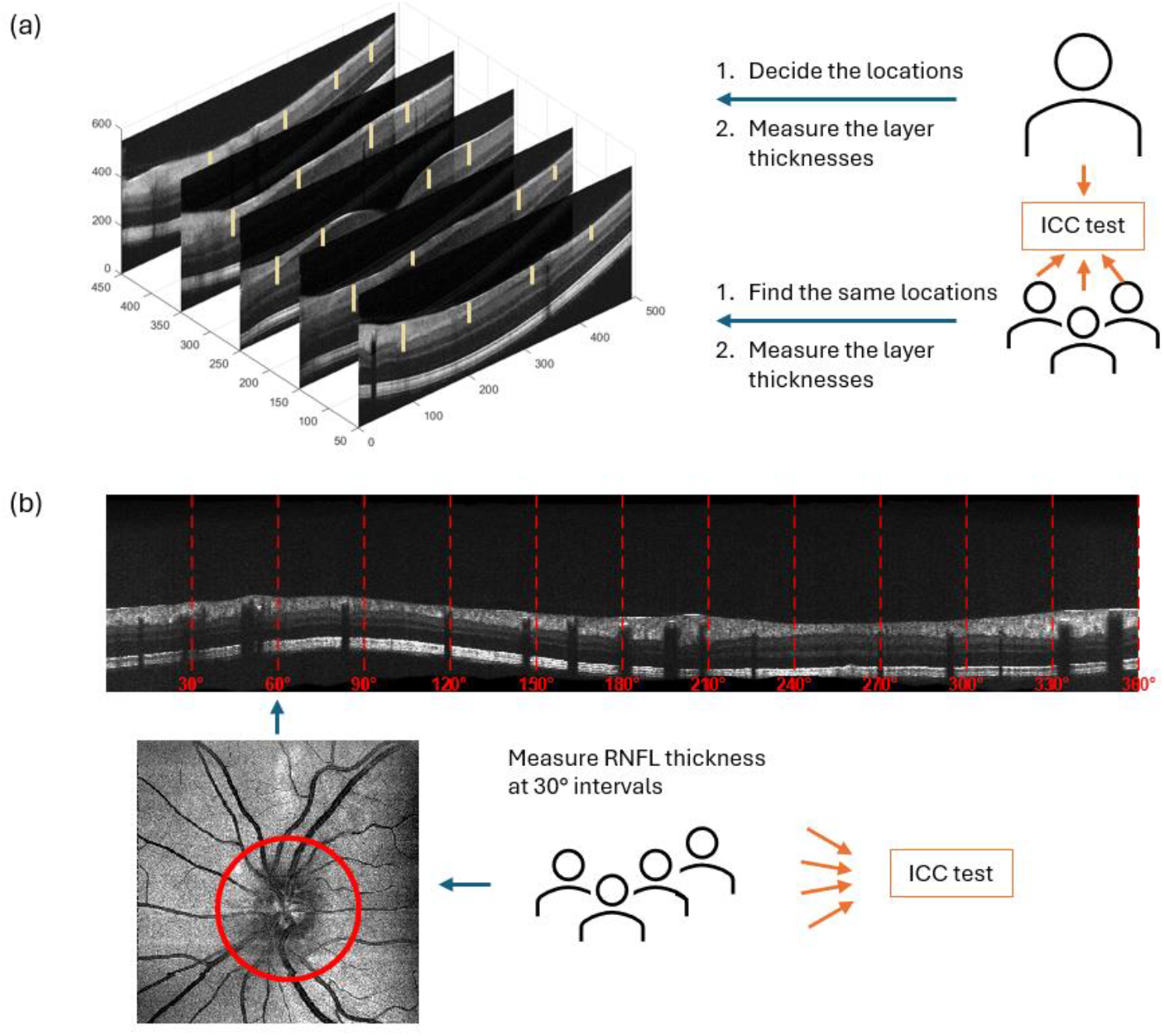
Repeatability test involving five operators on (a) macular-centered and (b) ONH-centered images.

To assess inter-operator consistency, we calculated the intraclass correlation coefficient (ICC) and used the ICC(3,1) model [25], a two-way mixed-effects model for single measurements with fixed raters. We selected this model to evaluate the reliability of individual raters’ measurements across all images. A high ICC(3,1) value (closer to 1) indicates strong inter-operator agreement, supporting the high quality and reproducibility of the Vis-OCT Explorer reconstructed images.

## Results

### 3.1 Validation of vis-OCT image reconstruction methods

Figure 4 shows that k-linearization, optimized dispersion compensation, and balanced detection substantially improved the reconstructed vis-OCT image quality. Fig. 4a is the direct FFT result from the raw data, and Figs. 4b&4c are the enlarged regions in the inner and outer retinal regions. With k-linearization in the raw data, major structural features are revealed after FFT, as shown in Figs. 4d-4f. After optimized dispersion compensation, sublayers of the outer retina became visible in the reconstructed images, as shown in Figs. 4g-4i. Applying balanced detection further suppressed background noise and enhanced differentiation between the ganglion cell layer (GCL) and the inner plexiform layer (IPL) (Figs. 4j-4l). These improvements enable visualization of fine retinal structures that are not easily observed with near-infrared OCT or unoptimized reconstructions, potentially providing additional information for disease diagnosis and intervention.

**Figure 4.**
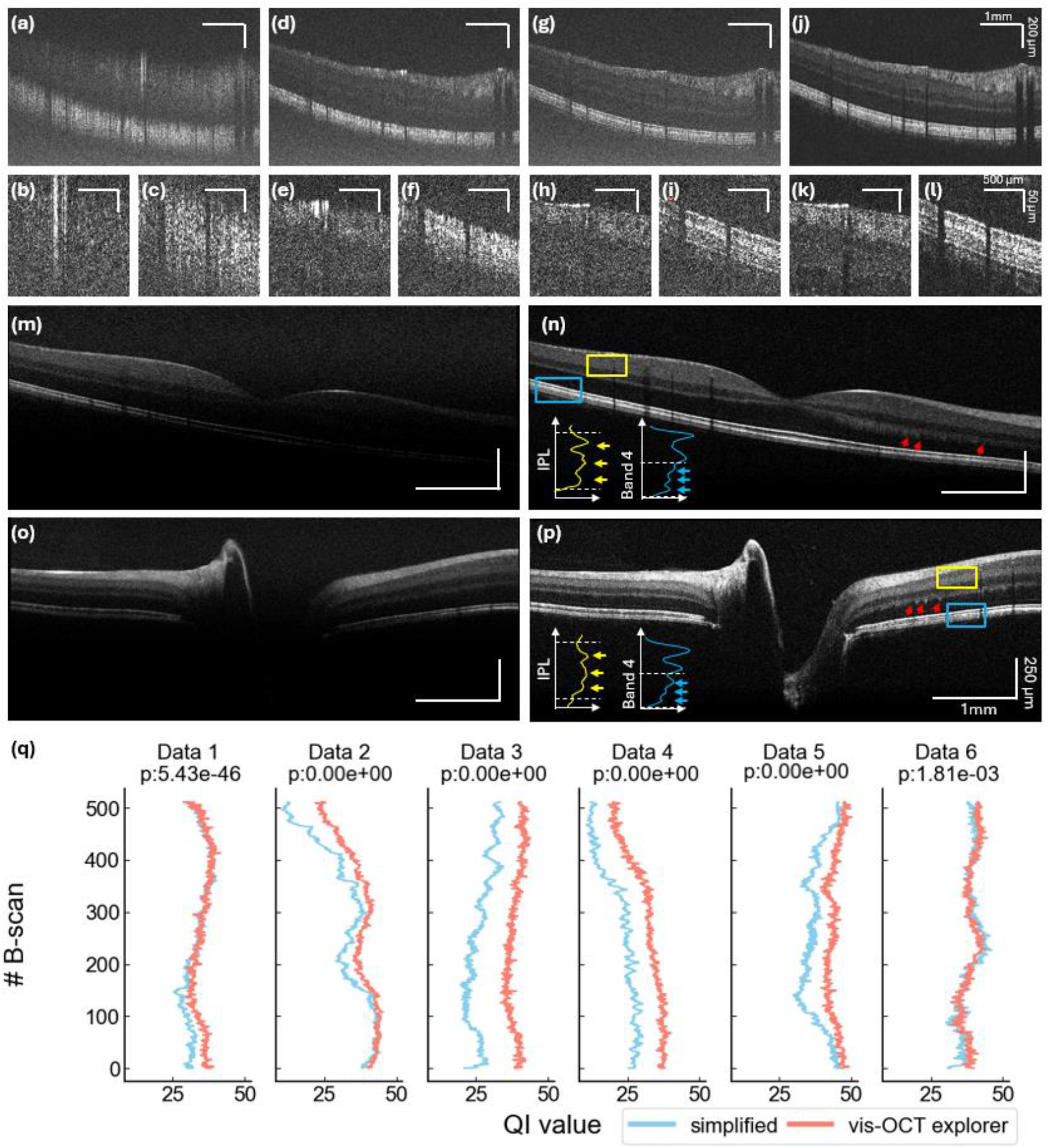
Comparison of reconstructed image quality with (a-c) direct FFT, (d-f) k-linearization + FFT, (g-i) k-linearization + dispersion compensation + FFT, and (j-l) balanced detection + k-linearization + dispersion compensation + FFT. (a), (d), (g), (j) show entire B-scans, with (b), (e), (h), and (k) showing magnified inner retinal regions, and (c), (f), (i), and (l) showing magnified outer retinal regions. A comparison between the (m) and (o) traditional reconstruction pipeline results and (n) and (p) Vis-OCT Explorer reconstructed images are presented. The red arrows point to the Henley fibers. The yellow and blue boxes highlight the sublayers of IPL and band 4 [26], respectively, with the line profile in each box shown on the side in the corresponding color. The yellow and blue arrows point to the sublayers in the IPL and band IV. (q) shows QI comparison between traditional processing (simplified, blue) and Vis-OCT Explorer processing (red) across B-scans on six datasets.

Figs. 4m–4p compared traditional OCT image reconstruction pipeline results with Vis-OCT Explorer processing for both macular (Figs. 4m&n) and ONH-centered (Figs. 4o&p) scans. All images were averaged using 10 neighboring B-scans to reduce speckle noise. Traditional processing results are shown on the left, where linear interpolation for balanced detection resulted in greater signal roll-off. In contrast, the Vis-OCT Explorer processing shown on the right revealed refined retinal layer structures, including IPL sublayers on the macular images (as highlighted by the yellow boxes), Henle’s fiber layer near the macula and ONH (as highlighted by the red arrows), and the clear separation between RPE and Bruch’s membrane (as highlighted by the blue boxes). For better visualization of sublayers, the intensity profiles of the segments in the yellow boxes are shown as the yellow lines in Figs 4n&p, where the yellow arrows point to the three peak borders separating ON and OFF sublamellae in the IPL [10]. The intensity profiles of segments in the blue boxes are shown as the blue lines in Figs 4n&p, where the blue arrows highlight the three hyperreflective bands in band IV, corresponding to rod outer segment tips, RPE, and inner boundary of the BM [26].

Fig. 4q presents the quantitative comparison results. We calculated the QI for each of the 512 B-scans from six balanced-detection human datasets reconstructed with the traditional processing method (blue) and Vis-OCT Explorer (red). P-values from paired Student’s t-tests were reported for each dataset, with p < 0.05 indicating significant improvement in image quality using the optimized reconstruction approach.

The Vis-OCT Explorer processing pipeline is also compatible with small animal datasets, enabling high-quality reconstructions suitable for biological research. Figs. 5a&5b show representative structural OCT and OCTA en face images of a mouse retina with the ONH positioned at the lower right of the FOV. In the structural OCT image, the black curving lines are shadows created by superficial retinal vessels, while black dots mark the positions of deep vessels without revealing their full extent. The OCTA *en-face* image delineates the whole network of superficial vascular plexus and deep capillary plexus as bright signals. Figs. 5c&5d show a tree shrew retina imaged approximately 1 mm from the ONH. Unlike the mouse *en face* image, where retinal ganglion cell axon bundles are not visible, the structural OCT *en-face* image of the tree shrew clearly highlights these axon bundles as tightly packed white stripes arranged in an orderly pattern between vessels, a configuration consistent with the tree shrew’s high visual acuity [18] The corresponding OCTA image, acquired in this peripheral region, reveals a dense deep capillary network as bright signals.

**Figure 5.**
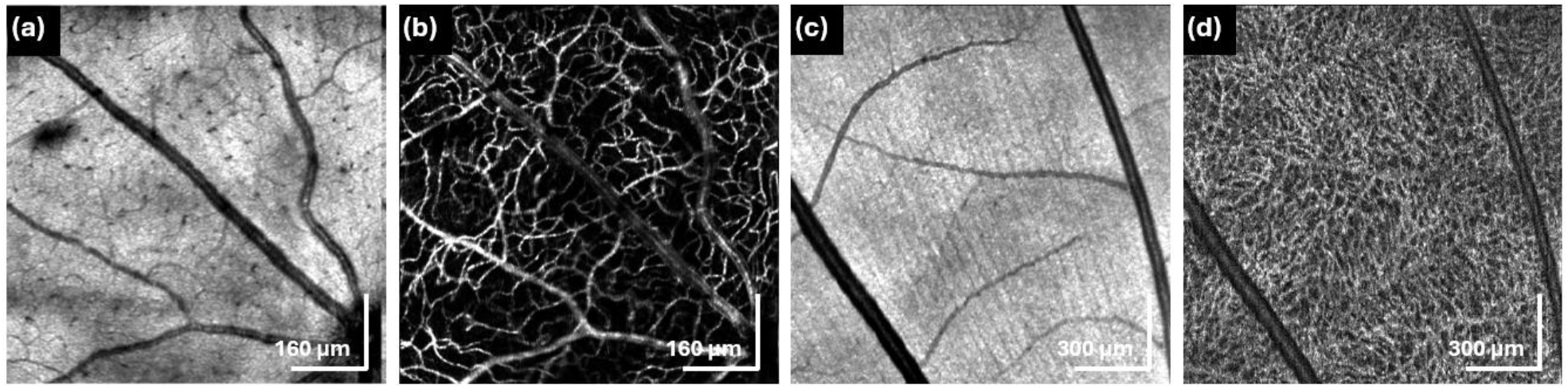
Representative reconstructed *en fac*e images from small animal datasets: (a) mouse structural OCT, (b) mouse OCTA, (c) tree shrew structural OCT, and (d) tree shrew OCTA.

### 3.2 Repeatability of image measurements

Figure 6 shows ICC(3,1) values of the repeated measurement tests for each layer measurement on each of the eight selected human datasets. On the macular images, the RNFL layer thickness measurement reproducibility reaches excellent and good levels [18]. Despite the IPL thickness measurement’s relatively low consistency, most of the GCL or GCIPL layer measurements result in excellent and good consistency. The deficiency in IPL measurements might be attributed to the sublayers existing in the IPL, which makes it hard to manually identify the true IPL boundaries. On the circular resampled B-scan around the ONH, the RNFL layer thickness measurements also achieve good consistency.

**Figure 6.**
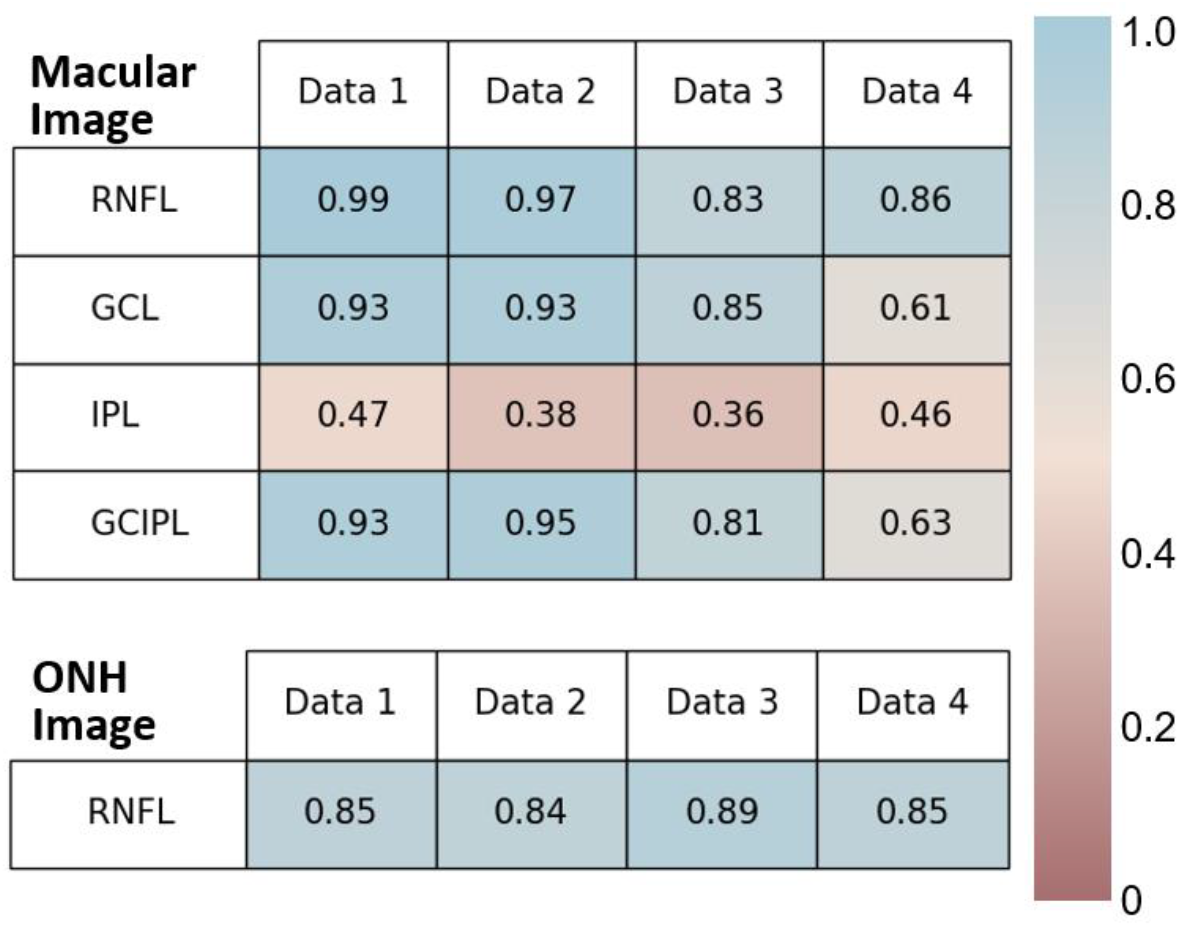
Repeatability test showing the ICC3 values for different layer measurements on human datasets.

### 3.3 Vis-OCT Explorer processing time consumption

We investigated the time usages for reconstructing vis-OCT volumes with Vis-OCT Explorer. Processing a structural vis-OCT volume containing 512 A-lines per B-scan and 512 B-scans took 360 seconds using the CPU version. With GPU acceleration, we reduced the processing time to 76 seconds using the NVIDIA GeForce RTX 3090 and 45 seconds using the NVIDIA GeForce RTX 4090 GPU. Additionally, reconstructing an OCTA volume of the same sampling rate with two repeated B-scans at each position and two repeated volumes, including registration and averaging, took approximately 800 seconds using the CPU version. With GPU acceleration, the entire process took 104 seconds using the NVIDIA GeForce RTX 3090 and 77 seconds using the NVIDIA GeForce RTX 4090.

## 4. Discussion

### 4.1 Vis-OCT Explorer for clinical research

High-quality post-processing is crucial for OCT, particularly vis-OCT, which offers higher axial resolution and unique contrast compared to near-infrared OCT, enabling more detailed visualization of retinal layers. For example, vis-OCT provides a clear contrast between the GCL and IPL, allowing for easy differentiation between these two layers. Its sub-2 µm axial resolution reveals three hyper-reflective and two hypo-reflective bands within the IPL, each 10–22 microns thick. While the thickness of the GCL and IPL complex (GCIPL) is commonly used for glaucoma diagnosis, evidence suggests that ganglion cell damage begins with their synapses in the IPL [27], and IPL thickness correlates strongly with glaucoma severity [28]. However, near-infrared OCT lacks the contrast and resolution to capture IPL details [29]. On vis-OCT, extracting these features depends on precise balanced detection [9] and dispersion compensation [10]. Vis-OCT Explorer provides open-source tools for this level of high-quality processing, enabling researchers to obtain detailed in vivo imaging information and advancing biological and clinical research.

The measurement repeatability test confirmed the functionality of the developed software. Clinical biomarkers such as RNFL and GCIPL thickness measurements showed good repeatability across different operators. Although the IPL measurement test was less consistent due to the lack of optimized manual labeling tools, we will improve IPL measurement in future updates. For example, current measurements were performed at a single A-line position, but extending the IPL labeling to all A-lines in a B-scan could incorporate adjacent information and potentially improve IPL measurement accuracy. Additionally, integrating automatic retinal layer thickness segmentation, based on edge detection or artificial intelligence [30-32], into the Vis-OCT Explorer will help alleviate manual identification and yield more consistent results. Future work will also compare Vis-OCT Explorer measurements with commercial software to assess consistency across devices for the same patient.

### 4.2 Vis-OCT Explorer for non-clinical research

We provide algorithms and code for the OCT image post-processing and reconstruction from raw fringe data. Raw fringe data carries additional information on the retina and other tissue changes compared to the reconstructed images. For instance, phase variations in the OCT interference pattern reveal nanoscale optical path length changes, enabling studies on retinal blood flow [33, 34], tissue elasticity [35, 36], and cone photoreceptor length changes [37]. Additionally, the spectral analysis of OCT raw fringe data reveals invaluable information from *in vivo* imaging, particularly in vis-OCT. The absorption spectra of melanin, macular pigment, and hemoglobin in the visible light range provide critical insights for diagnosing outer retinal diseases and calculating retinal blood oxygenation [38-41].

However, most commercial OCT systems do not provide access to raw fringe data, and no OCT database of spectral domain data currently exists. This limitation partially stems from the lack of widely shared algorithms for processing raw fringe data. In contrast, the MRI community has established databases containing both k-space data and reconstructed DICOM images [42, 43], supported by open-source reconstruction software and AI-based developments [44]. The National Eye Institute’s (NEI) recent notice (NOT-EY-24-006) emphasizes access to raw data in the ocular imaging community. Sharing raw interferogram data could significantly advance OCT image reconstruction research, attracting contributions from the OCT community, computer scientists, and engineers. Therefore, Vis-OCT Explorer can set an important precedent for open-source software for processing raw interferogram data and promote standardization across the field.

This platform enables non-clinical researchers to engage in advanced image post-processing. The software packages were developed in the Python environment, where abundant computer vision and neural network-based image processing packages are available. Further development could expand the functions of Vis-OCT Explorer. For example, retinal layer segmentation and disease classification can be potentially integrated into the current software with OpenCV, TensorFlow, SciPy, and any future Python packages. Additional phase and spectral analyses of the raw data could also be integrated in the future. Despite the software being developed based on vis-OCT datasets, with minimal modifications, similar algorithms could be applied to near-infrared OCT datasets.

## Conclusion

We present Vis-OCT Explorer, an open-source toolbox for high-quality vis-OCT image reconstruction, freely available to the OCT community. It addresses the OCT post-processing challenges by providing tools for balanced detection, dispersion compensation, and enhanced image reconstruction, enabling detailed visualization of retinal structures. The software demonstrates repeatability on commonly used retinal thickness measurements in the clinics and provides the option to process animal data. In this way, Vis-OCT Explorer bridges the gap between raw data and clinically meaningful outputs. By building on platforms like this, future researchers could develop advanced post-processing techniques, making raw fringe data more accessible and paving the way for data standardization across the field.

## Credit authorship contribution statement

**Weijia Fan:** Writing – original draft, Validation, Methodology, Investigation, Data curation, Formal analysis, Visualization. **Fengyuanshan Xu:** Writing – review & editing, Validation, Software, Resources, Methodology, Investigation, Formal analysis. **Roman Kuranov:** Investigation, Methodology. **Ronald Zambrano, Jacqueline Chen, Jiahui Wu, Seung Lee, Kenny Q. Trang, Rukhsana Mirza, Shira Simon, Fabio Lavinsky, Alex S. Huang, Xiaorong Liu, Jeffrey L. Goldberg, Joel S. Schuman:** Investigation, Validation. **Hao F. Zhang:** Investigation, Validation, Writing– review & editing, Funding acquisition, Supervision

